# Lack of evidence for microbiota in the placental and fetal tissues of rhesus macaques

**DOI:** 10.1101/2020.03.05.980052

**Authors:** Kevin R. Theis, Roberto Romero, Andrew D. Winters, Alan H. Jobe, Nardhy Gomez-Lopez

## Abstract

The prevailing paradigm in obstetrics has been the sterile womb hypothesis. However, some are asserting that the placenta, intra-amniotic environment, and fetus harbor microbial communities. The objective of this study was to determine if the fetal and placental tissues of rhesus macaques harbor viable bacterial communities. Fetal, placental, and uterine wall samples were obtained from cesarean deliveries without labor (∼130/166 days gestation). The presence of viable bacteria in the fetal intestine and placenta was investigated through culture. The bacterial burden and profile of the placenta, umbilical cord, and fetal brain, heart, liver, and colon were determined through quantitative real-time PCR and DNA sequencing. These data were compared with those of the uterine wall, as well as to negative and positive technical controls. Bacterial cultures of fetal and placental tissues yielded only a single colony of *Cutibacterium acnes*. This bacterium was detected at a low relative abundance (0.02%) in the 16S rRNA gene profile of the villous tree sample from which it was cultured, yet it was also identified in 12/29 background technical controls. The bacterial burden and profile of fetal and placental tissues did not exceed or differ from those of background technical controls. In contrast, the bacterial burden and profiles of positive controls exceeded and differed from those of background controls. Among the macaque samples, distinct microbial signals were limited to the uterine wall. Therefore, using multiple modes of microbiologic inquiry, there was not consistent evidence of viable bacterial communities in the fetal and placental tissues of rhesus macaques.

**IMPORTANCE:** Microbial invasion of the amniotic cavity (i.e. intra-amniotic infection) has been causally linked to pregnancy complications, especially preterm birth. Therefore, if the placenta and the fetus are typically populated by low biomass yet viable microbial communities, current understanding of the role of microbes in reproduction and pregnancy outcomes will need to be fundamentally reconsidered. Could these communities be of benefit by competitively excluding potential pathogens or priming the fetal immune system for the microbial bombardment it will experience upon delivery? If so, what properties (e.g. microbial load, community membership) of these microbial communities preclude versus promote intra-amniotic infection? Given the ramifications of the *in utero* colonization hypothesis, critical evaluation is required. In this study, using multiple modes of microbiologic inquiry (i.e. culture, qPCR, DNA sequencing) and controlling for potential background DNA contamination, we did not find consistent evidence for microbial communities in the placenta and fetal tissues of rhesus macaques.

## INTRODUCTION

The development and widespread use of DNA sequencing technologies to characterize host-associated microbial communities has increasingly led researchers to question the sterility of body sites and fluids previously presumed to be free of resident microorganisms. For example, researchers have recently proposed the existence of microbiota in the human blood (1–8), bladder (9–16), uterus (17–30), placenta (31–45), and fetus (36, 44–46). This has led to discussion in the literature on the caveats associated with studies of the microbiota of very low microbial biomass, or potentially sterile, body sites (47–54). In particular, there has been much debate over the existence of a placental microbiota (31–45, 50, 55–69) and of *in utero* microbial colonization of the human fetus (36, 44–46, 64, 70–72).

The primary focus of the debate is that most of the studies proposing the existence of placental and fetal microbiota in humans have relied heavily, if not exclusively, on DNA sequencing techniques (31–35, 37–42, 45), and the bacterial signals in these studies may be background DNA contaminants from extraction kits, PCR and sequencing reagents, and general laboratory environments (50, 55, 57, 59, 62). Furthermore, even if the bacterial DNA sequence data are derived from placental and fetal tissues and not from background contamination, this does not necessarily indicate that there are viable bacterial communities in the placenta or the fetus. Specifically, the bacterial DNA sequence data may reflect bacterial products and components rather than resident microbiota (73–77).

As a consequence, we and others (50, 62) have suggested criteria for establishing the existence of placental and fetal microbiota. First, viability of the resident bacteria should be established through culture or metatranscriptomic data from bacterial-specific genes within placental and fetal tissues. Second, the bacterial load of placental and fetal tissues, as demonstrated through quantitative real-time PCR (qPCR), should exceed those of background technical controls. Third, the bacterial profiles of placental and fetal tissues should be distinct from those of the technical controls. Fourth, the resident bacteria should be visualized in the tissues through microscopy. Fifth, the taxonomic data of the detected bacteria should be ecologically plausible (50, 62). There have been many studies that may have met one or two of these criteria (31–46, 78–80), but no study has yet attempted to simultaneously meet all criteria and ultimately conclude that there is widespread colonization of the placenta and/or fetus by viable microbial communities (72).

Although most of the research evaluating the existence of placental and fetal microbiota has been done with human subjects, animal models afford opportunities to surgically obtain placental and fetal tissues before the process of labor. Tissues collected after the process of labor could confound experimental results regarding *in utero* colonization due to potential microbial invasion of the amniotic cavity (81–83). Several studies using rat and mouse models have provided mixed evidence: while three studies detected placental and fetal microbiota through DNA sequencing techniques following cesarean delivery (44, 46, 75), two other studies did not (60, 84). In non-human primates, specifically rhesus and Japanese macaques, a unique placental and/or fetal microbiota has been consistently detected through DNA sequencing following cesarean delivery (85–89). However, these preliminary studies neither include culture or qPCR components nor display the sequence data from background technical controls.

The objective of the current study was therefore to determine whether the fetal and placental tissues of rhesus macaques harbor bacterial communities using bacterial culture, qPCR, and 16S rRNA gene sequencing and by comparing the bacterial profiles of these tissues to those of background technical controls.

## RESULTS

### Bacterial culture from fetal and placental samples

All negative culture controls were negative (no bacterial growth over seven days) and all positive culture controls were positive (lawn of bacterial growth within 24 hours). The 96 total cultures of fetal and placental samples from the four rhesus macaques yielded only a single bacterial colony (**Figure 1**). This bacterium grew on an anaerobically incubated chocolate agar plate inoculated with the villous tree sample from Subject 1. A BLAST query of the 16S rRNA gene of this bacterium revealed that it was *Cutibacterium acnes*. Specifically, the 16S rRNA genes of this bacterium and American Type Culture Collection (ATCC) strain 6919 (Accession # NR_040847.1; *Cutibacterium acnes* Scholz and Kilian) were identical across 1,056 nucleotide bases.

**Figure 1.**
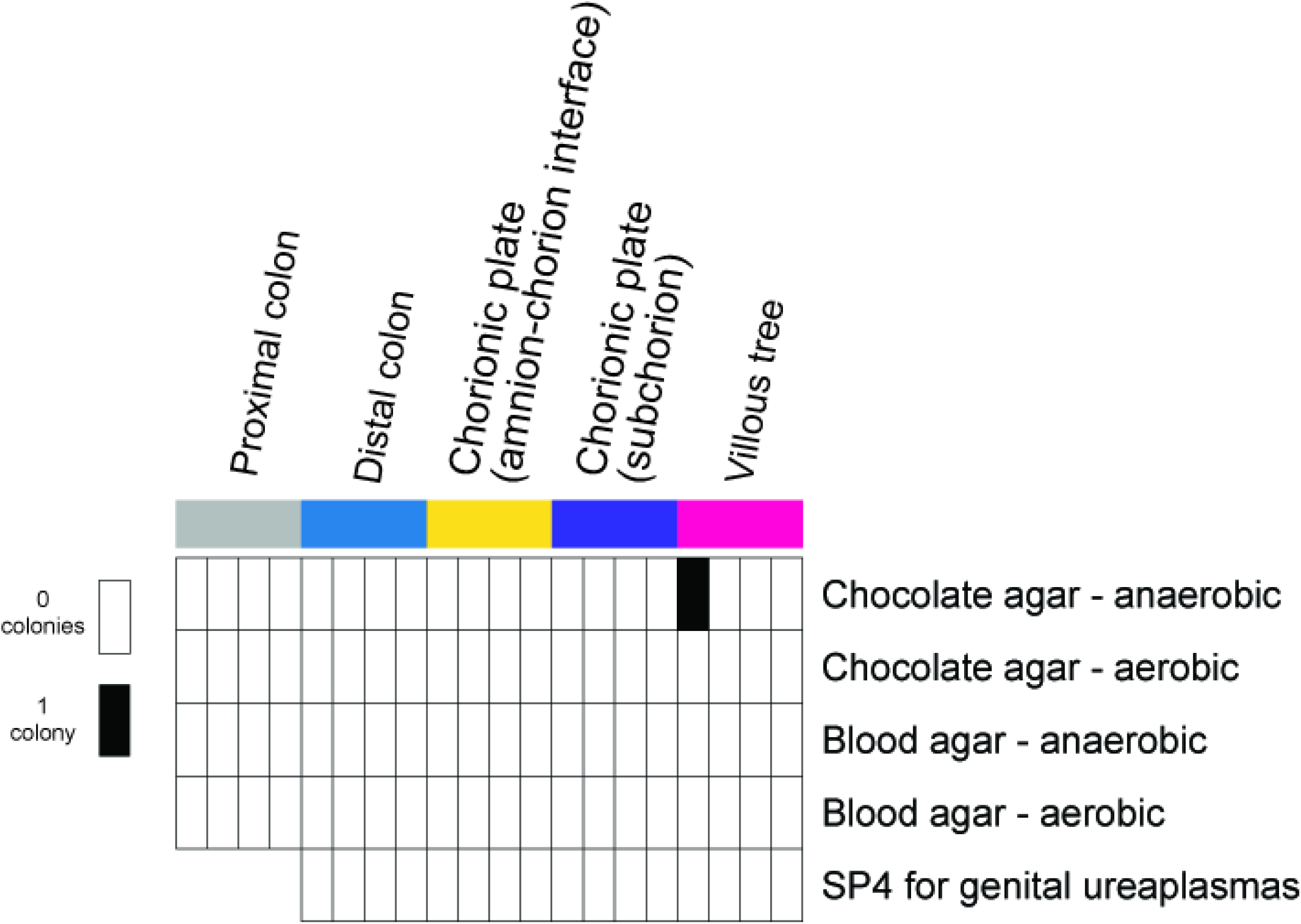
Results of bacterial culture of ESwabs of rhesus macaque fetal and placental tissues. ESwabs of the proximal colon, distal colon, chorionic plate (both amnion-chorion interface and the subchorion), and the villous tree were collected from each of the four subjects and plated on chocolate and blood agar, which was then incubated under anaerobic and aerobic conditions for seven days. SP4 broth was also inoculated to assess the presence of genital ureaplasmas.

It was next determined whether the 16S rRNA gene of this cultured bacterium was also present in the 16S rRNA gene profile of the villous tree sample from which it was recovered, as well as in the profiles of other fetal, placental, and uterine wall samples for Subject 1. The 16S rRNA gene profile of the villous tree swab sample from Subject 1 included 35,780 sequences and had a Good’s coverage value of 99.9% (i.e., the sample’s bacterial profile was thoroughly characterized). Seven of the 35,780 (0.02%) sequences from this sample were an exact match to the V4 region of the 16S rRNA gene of the cultured *Cutibacterium*. There was not an exact match to any of the 16S rRNA gene sequences in the bacterial profile of the villous tree and basal plate tissue (i.e. not a swab) sample for Subject 1, but this sample included only 78 sequences (i.e., it was not well characterized). Exact matches to the cultured *Cutibacterium*’s 16S rRNA gene were also identified in the bacterial profiles of the chorionic plate [chorionic plate tissue: 21/355 (5.9%) sequences; top of amnion swab: 1/87,899 (0.001%) sequences; amnion-chorion interface swab: 14/350 (4.0%) sequences], the umbilical cord [24/13,700 (0.18%) sequences], the fetal distal colon [286/106,663 (0.27%) sequences], and the fetal heart [163/11436 (1.4%) sequences] for Subject 1. Exact matches to the cultured *Cutibacterium*’s 16S rRNA gene were also identified in the bacterial profile of the decidua swab for Subject 1 [22,619/76,987 (29.4%) sequences].

Lastly, it was determined whether the 16S rRNA gene of this cultured bacterium was present in the 16S rRNA gene profiles (prior to subsampling) of the background technical controls. Exact matches to the 16S rRNA gene of this cultured *Cutibacterium* were identified in 5/14 (35.7%) sterile swab controls and 7/15 (46.7%) blank DNA extraction kit controls at average relative abundances of 0.46% (maximum 2.0%) and 4.55% (maximum 23.1%), respectively. Therefore, it is unclear if this *Cutibacterium* was present in fetal, placental, and uterine wall samples of Subject 1 or if it was a contaminant.

### Quantitative real-time PCR (qPCR) of fetal, placental, and uterine wall samples and controls

The bacterial burden of fetal and placental tissues did not exceed that of background technical controls (**Figure 2A**,**B**). Among the swab samples, only the maternal myometrium had a higher bacterial load than sterile swabs (Mann-Whitney U test: U = 0, p = 0.005; **Figure 2A**). No fetal, placental, or uterine wall tissue samples consistently had higher bacterial loads than blank DNA extraction kits (**Figure 2B**).

**Figure 2.**
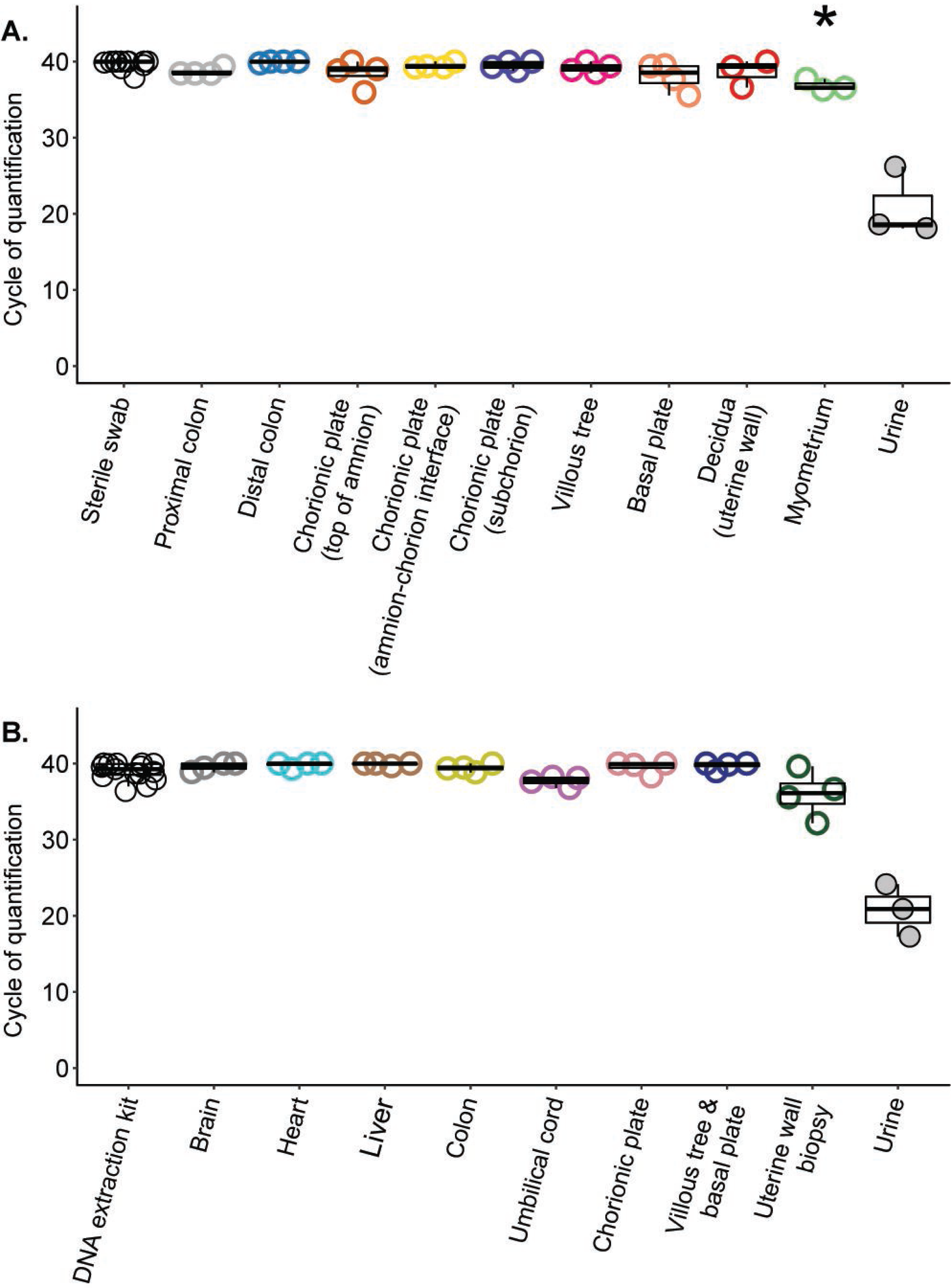
Quantitative real-time PCR (qPCR) analyses illustrating the cycle of quantification values among rhesus macaque fetal, placental, and uterine wall A) swab and B) tissue samples, and their respective negative technical controls. The negative controls for swab and tissue samples were DNA extraction kits processed with and without a sterile Dacron swab, respectively. The positive controls are human urine samples. In the plots, lower cycle of quantification values indicate higher bacterial loads. Bars indicate the median and quartile cycle of quantification values for each sample and control type. Points, color-coded by sample type, indicate the mean values of three replicate qPCR reactions. An asterisk indicates that bacterial loads of that swab or tissue sample type were greater than those of corresponding negative technical controls based on Mann-Whitney U or t-tests with sequential Bonferroni corrections applied.

### 16S rRNA gene sequencing of fetal, placental, and uterine wall samples and controls

Twelve of the 14 (85.7%) sterile swab controls and 10/15 (66.7%) blank DNA extraction kits yielded a 16S rRNA gene library with ≥ 500 quality-filtered sequences and a Good’s coverage ≥ 95%. Twenty-six of 28 (92.9%) fetal and placental swab samples and 22/28 (78.6%) fetal and placental tissue samples yielded 16S rRNA gene libraries meeting these criteria, as did all (10/10) uterine wall samples and all (3/3) human urine positive controls. These samples were included in 16S rRNA gene profile analyses.

With respect to alpha diversity, there were no swab or tissue sample types from rhesus macaques whose amplicon sequence variant (ASV) profiles had a richness (i.e. Chao1 index) or heterogeneity (i.e. Shannon and Simpson indices) that differed from those of their respective negative technical control. Human urine samples also did not have ASV profiles that differed in richness or heterogeneity from the sterile swab controls.

With respect to beta diversity, the overall ASV profiles of fetal and placental swab and tissue samples did not differ from those of their respective technical control (NPMANOVA using the Bray-Curtis similarity index; p ≥ 0.21; **Figure 3**). The ASV profiles of swabs of the myometrium (F = 1.739, p = 0.0094), but not the decidua (F = 0.9193, p = 0.64), differed from the profiles of sterile swabs (**Figure 3A**). Similarly, the ASV profiles of uterine wall biopsies differed from those of blank DNA extraction kits (F = 1.860, p = 0.0076; **Figure 3B**). The ASV profiles of human urine also differed from those of sterile swab controls (F = 1.834, p = 0.0058).

**Figure 3.**
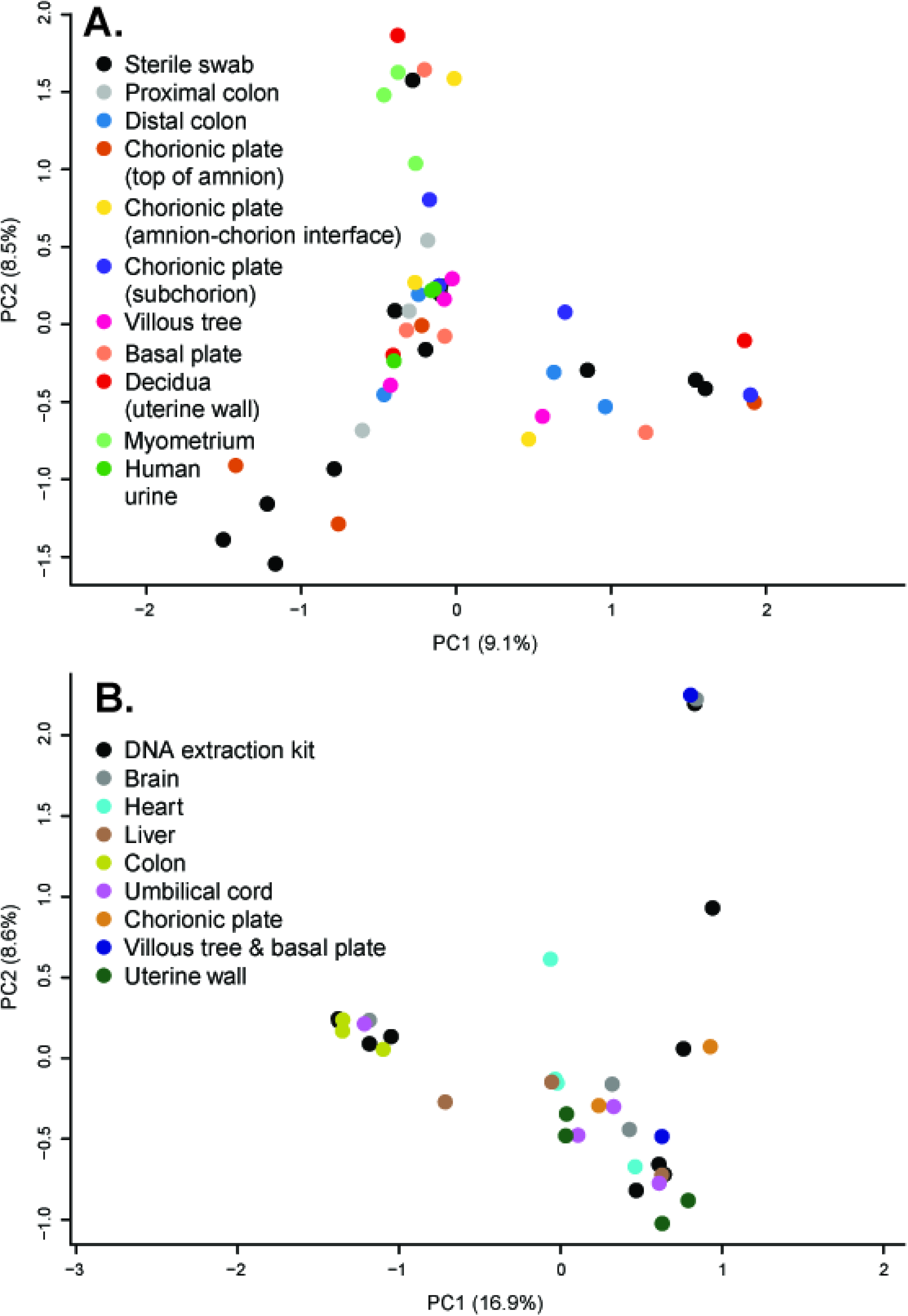
Principal Coordinates Analyses (PCoA) illustrating variation in 16S rRNA gene profiles among fetal, placental, and uterine wall A) swab and B) tissue samples, and their respective negative technical controls. 16S rRNA gene profiles were characterized using the Bray-Curtis similarity index.

The bacterial taxonomic data associated with the ASV profiles of the fetal, placental, and uterine wall samples and controls are illustrated in **Figure 4**. There were only two prominent (≥ 5% relative abundance) ASVs among the fetal and placental swab and tissue samples: ASVs 001 (*Staphylococcus*) and 002 (*Pelomonas*). These two ASVs were also prominent in the profiles of both the sterile swabs and the blank DNA extraction kits. ASV 001 was identified in the profiles of 9/12 (75%) and 5/10 (50%) swab and extraction kit technical controls, respectively, while ASV 002 was identified in 5/12 (42%) sterile swab and 6/10 (60%) extraction kit profiles. ASV 001, but not ASV 002, was identified as a contaminant among swab samples by the decontam program (**Figure 4**). Among tissue samples, neither ASV 001 or ASV 002 were identified as contaminants using decontam.

**Figure 4.**
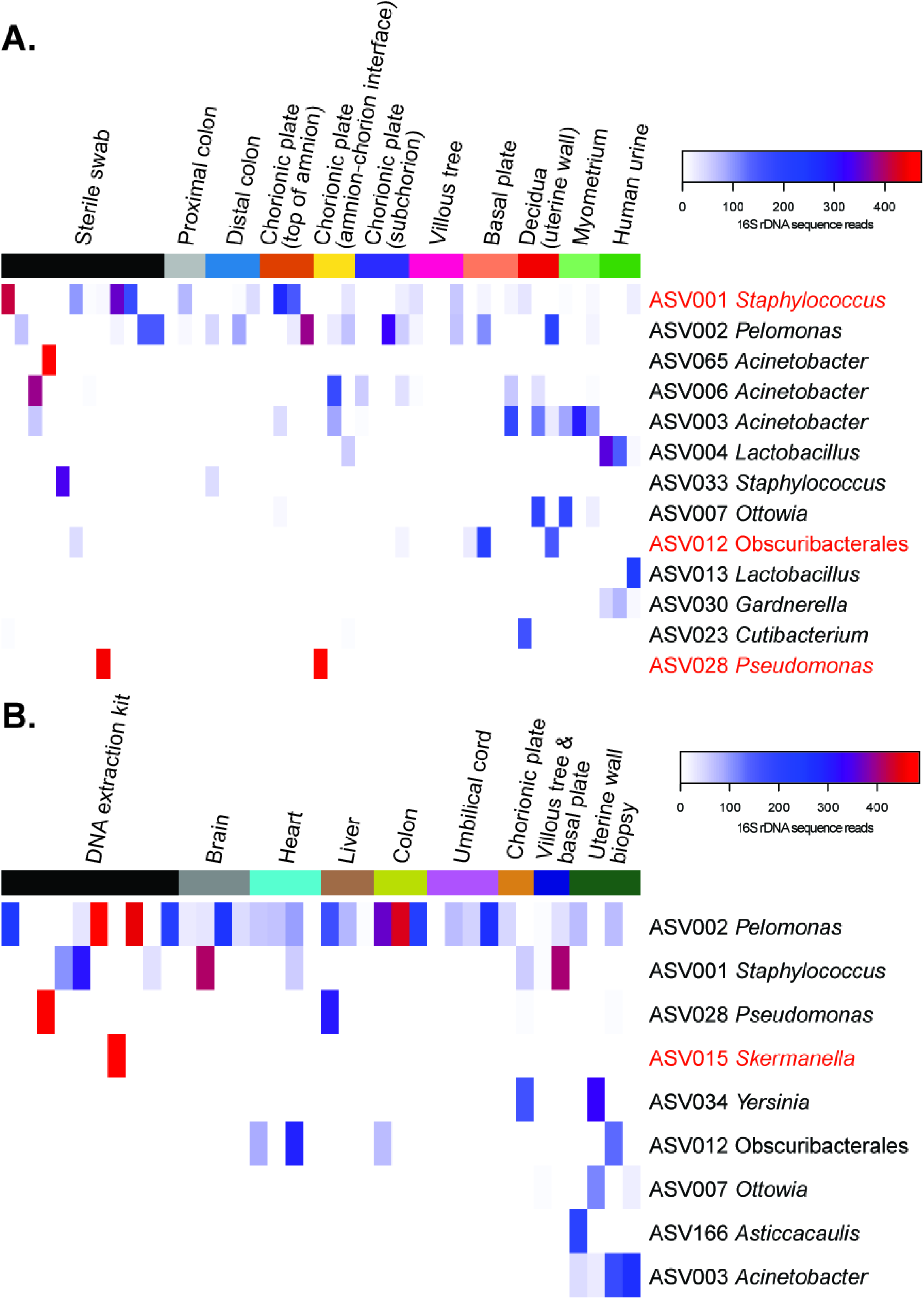
Heat map illustrating the relative abundances of prominent (≥ 5% average relative abundance) amplicon sequence variants (ASVs) among the 16S rRNA gene profiles of fetal, placental, and uterine wall A) swab and B) tissue samples, and their respective negative technical controls. Human urine samples are included as positive technical controls in panel A. The four ASVs in red font were identified as background DNA contaminants by the R package decontam.

Aside from ASV 002, ASVs 003 (*Acinetobacter*), 007 (*Ottowia*), and 012 (uncl. Obscuribacterales) were prominent (≥ 5% relative abundance) among both uterine wall swab and tissue samples (**Figure 4**). None of these three ASVs were prominent among either sterile swab or extraction kit technical controls, but ASV 012 (uncl. Obscuribacterales) was identified as a contaminant by the decontam program among the swab samples. ASV 003 (*Acinetobacter*) was identified in 2/3 decidua swab, 3/3 myometrium swab, and 4/4 uterine wall biopsy samples, with an average relative abundance of 22.9%. Conversely, it was identified in only 1/12 sterile swab and 0/10 blank extraction kit controls. ASV 007 (*Ottowia*) was identified in 1/3 decidua swab, 2/3 myometrium swab, and 3/4 uterine wall biopsy samples, with an average relative abundance of 10.5%. This ASV was not identified in any sterile swab or blank extraction kit control.

Human urine samples were sequenced alongside the rhesus macaque swab samples to serve as low microbial biomass positive controls. The prominent (≥ 5% relative abundance) ASVs among the urine samples were 004 (*Lactobacillus*), 013 (*Lactobacillus*), and 030 (*Gardnerella*). These three ASVs were not identified in the profiles of any of the sterile swabs.

## DISCUSSION

### Principal findings of the study

First, recovery of bacterial cultures from the fetal and placental tissues of rhesus macaques was very rare. The 96 cultures performed yielded only a single colony of *Cutibacterium* (*Propionibacterium*) *acnes*. Second, the bacterial burden of fetal and placental samples did not exceed that of background technical controls. Third, the bacterial profiles of fetal and placental samples did not differ from those of background technical controls. Fourth, among the intrauterine sites of the rhesus macaque investigated here, only the uterine wall exhibited a distinct microbial signature.

### Prior reports of fetal and placental microbiota in non-human primates

There have been five preliminary studies (85–89) of fetal (130-139 days gestation) and/or placental microbiota in rhesus and Japanese macaques and the collective conclusion was that the intrauterine environment, the fetus, and the placenta were colonized by bacterial communities. In the first three studies, rhesus or Japanese macaque dams received control or high fat diets. In the first study (85), the bacterial profiles of the fetal colon and oral cavity of rhesus macaques were compared to those of the placenta and the maternal anal, vaginal, and oral cavity using 16S rRNA gene sequencing. The bacterial profiles of fetal samples were similar to those of the placenta but distinct from those of maternal samples. The bacteria reportedly inhabiting the fetus (i.e. *Acinetobacter, Propionibacterium, Streptococcus, Staphylococcus*, and *Bacteroides*) appeared to be derived from the placental microbiota (85). In the second study (86), the bacterial profiles of the fetal colon and oral cavity of Japanese macaques were compared between dams receiving control and high-fat diets using 16S rRNA gene sequencing. The fetal bacterial profiles differed between the two treatment groups and the bacterial profiles of the offspring of dams receiving a high-fat diet exhibited a higher relative abundance of Pasteurellaceae than did the profiles of offspring from control dams. In the third study (87), the bacterial profiles of the fetal colon of Japanese macaques were compared with those of the developing infant colon at six and 10 months of age using 16S rRNA gene and shotgun metagenomic sequencing. Predominant members of the infant gut microbiota were often identified in the bacterial profiles of the fetal colon.

In the fourth and fifth studies, rhesus macaque dams received intra-amniotic injections of saline, lipopolysaccharide, interleukin 1 β, or *Ureaplasma parvum* to serve as a primate model of inflammatory preterm birth. In the fourth study (88), the intra-amniotic injection of inflammatory inducers (i.e. lipopolysaccharide, interleukin 1 β, or *Ureaplasma parvum*) altered the bacterial profiles of the placenta. In the fifth study (89), the intra-amniotic injection of inflammatory inducers again altered the bacterial profiles of the placenta; the bacterial profiles of placentas from control subjects exhibited a higher alpha diversity than those from subjects receiving inflammatory inducers. Relatively abundant taxa within the placental bacterial profiles of control subjects included *Acinetobacter, Agrobacterium, Bacteroides, Blautia, Cloacibacterium, Faecalibacterium, Haemophilus, Lactobacillus, Oscillospira, Porphyromonas, Prevotella*, and *Streptococcus*.

These five preliminary studies (85–89) provided initial investigations into the existence of fetal and/or placental microbiota in non-human primates. However, these preliminary studies did not include culture or qPCR components and, although DNA extraction and sequencing controls were mentioned in descriptions of the study design, the data from these controls were not presented or incorporated into the analyses of the bacterial profiles of fetal and placental samples. Therefore, it is unknown if the reported bacterial signals were distinct from or greater than those present in background technical controls. Even if the bacterial signals from fetal and placental samples were distinct from those in controls, it is still unknown if they are derived from viable microbiota inhabiting the fetal and placental compartments of macaques.

### The findings of this study in the context of prior reports

The current study includes culture and qPCR components and incorporates data from background technical controls into the analysis and evaluation of the existence of fetal, placental and uterine wall microbiota.

The collective bacterial cultures in this study yielded only a single isolate; one colony of *Cutibacterium* (*Propionibacterium*) *acnes* was obtained from a villous tree sample. The 16S rRNA gene of this bacterium was identified in the molecular surveys of this villous tree sample, as well as in the molecular surveys of the chorionic plate, umbilical cord, fetal colon and fetal heart samples from this subject. The relative abundance of this bacterium in the 16S rRNA gene profile of the villous tree sample was very low (0.02%), but its relative abundance in the swab of the maternal decidua sample from this subject was 29.4%. Given that this bacterium was cultured from the villous tree, was identified in molecular surveys of the villous tree sample and other placental and fetal samples from this subject, and was further detected at high relative abundances in a maternal decidua sample for this subject, it is reasonable to consider whether this isolate represents a viable bacterium that was transmitted from the mother to the fetus through the placenta. *Cutibacterium* (*Propionibacterium*) *acnes* has also been cultured from the human placenta and intra-amniotic environment. For instance, in a recent study concluding there exists distinct microbial communities in the human placenta and amniotic fluid in normal term pregnancies (36), 17/24 (70.8%) bacterial isolates obtained from placental tissues and amniotic fluids were *Propionibacterium* spp. and 5/24 (20.8%) were specifically *Cutibacterium* (*Propionibacterium*) *acnes*. However, in the current study, the 16S rRNA gene of the cultured *Cutibacterium* (*Propionibacterium*) *acnes* was identified in the molecular surveys of 12/29 (41.4%) background technical controls and in the bacterial profile of one blank DNA extraction kit this 16S rRNA gene variant constituted 23.1% of the sequences. Furthermore, *Cutibacterium* (*Propionibacterium*) *acnes* is a typical member of the human skin microbiota (90). Therefore, it is also reasonable to consider whether this isolate and molecular signals of *Cutibacterium*/*Propionibacterium* may simply represent microbial contamination from study personnel.

In the current study, qPCR revealed that the quantities of 16S rRNA gene copies in the placenta (i.e. basal plate, villous tree, and the subchorion, amnion-chorion interface, and amnion of the chorionic plate), umbilical cord, and fetal organs (i.e. brain, heart, liver, colon) of rhesus macaques did not exceed those in background technical controls (i.e. sterile swabs and DNA extraction kits). These results are consistent with those of prior studies showing that the quantities of 16S rRNA gene copies in the human placenta are indistinguishable from those of background technical controls (55, 57, 62).

In this study, there were no fetal or placental sites whose 16S rRNA gene profiles differed from those of background technical controls. Among the fetal and placental samples there were only two prominent (i.e. ≥ 5% average relative abundance) ASVs – they were classified as *Staphylococcus* and *Pelomonas. Staphylococcus* is a genus of bacteria commonly associated with mammalian skin and mucosal surfaces (91). It was detected in a preliminary molecular survey of the fetus and placenta of the Japanese macaque (85) and it has also been reported in numerous DNA sequence-based investigations of the human placenta (31–38, 41–43). However, *Staphylococcus* has also been identified as a background DNA contaminant in sequence-based studies (48), including in several prior studies of the human placenta (39, 57, 62). In the current study, the prominent ASV classified as *Staphylococcus* was also prominent and widespread among the background technical control samples, suggesting that it was a background DNA contaminant in this study as well.

*Pelomonas* is a genus of bacteria previously isolated from mud, industrial water, and hemodialysis water (92, 93). *Pelomonas* was not reported in prior preliminary molecular surveys of the fetal and placental tissues of macaques (85, 89), and it has only been reported in a single study as being a component of the human placental microbiota (41). Yet, *Pelomonas* has been identified as a background DNA contaminant in sequence-based studies (47, 48, 94), including in prior studies of the human placenta (57, 59, 62). As with *Staphylococcus*, in the current study, the prominent ASV classified as *Pelomonas* was also prominent and widespread among the background technical control samples, suggesting that it was a background DNA contaminant.

The only rhesus macaque samples with bacterial profiles distinct from those of background technical controls were the myometrial swabs and the uterine wall biopsies. These sample types also had the highest bacterial load, as assessed through qPCR. There were four ASVs that were prominent (i.e. ≥ 5% average relative abundance) among all uterine wall samples – they were classified as *Acinetobacter, Ottowia, Pelomonas*, and a member of the order Obscuribacterales. As discussed above, the ASV classified as *Pelomonas* is likely a DNA contaminant. Also, the program decontam identified the ASV classified as Obscuribacterales as another likely DNA contaminant. The data from *Acinetobacter* and *Ottowia* are more compelling. The primary ASV classified as *Acinetobacter* was detected in 9/10 (90%) uterine wall samples at an average relative abundance of 22.9%. In contrast, it was detected in only 1/22 (4.5%) background technical controls. *Acinetobacter* has been reported in prior sequence-based investigations of the human endometrium (20, 23, 24, 29, 30), and it has been cultured from the human endometrium as well (95). The primary ASV classified as *Ottawia* was detected in 6/10 (60%) uterine wall samples at an average relative abundance of 10.5%. It was not detected in any background technical controls. *Ottowia* is a genus of bacteria that has been isolated from industrial and municipal wastewater (96–99), sikhye (100), tofu residue (101), and fish intestines (102); it has not been identified in investigations of the human uterus. Nevertheless, *Ottowia* is a member of the family Comamonadaceae, and Chen et al (23) and Winters et al (30) reported that Comamonadaceae was among the most relatively abundant bacterial taxa in the human endometrium. Whether the molecular signals of *Acinetobacter* and *Ottowia* in the uterine wall in the current study represent a viable and residential uterine microbiota in rhesus macaques is unknown. However, the existence of uterine microbiota in non-human primates and the potential ramifications for female reproductive health warrant further investigation.

### Strengths of this study

First, this study included multiple modes of microbiologic inquiry, including bacterial culture, 16S rRNA gene qPCR, and 16S rRNA gene sequencing, to determine if the fetal, placental, and uterine wall tissues of rhesus macaques harbor bacterial communities. Second, placenta, fetal intestine, and uterine wall tissues were sampled both directly and through the use of swabs to enable verification of molecular microbiology results across sampling methods. Third, this study included low microbial biomass samples (i.e. human urine) to serve as technical positive controls for 16S rRNA gene qPCR and sequencing analyses. Fourth, controls for potential background DNA contamination were incorporated into 16S rRNA gene qPCR and sequencing analyses.

### Limitations of this study

First, given that the study was conducted on a non-human primate, the sample size was understandably low. Second, this study did not include fluorescent *in situ* hybridization or scanning electron microscopy to visualize potential microbial communities in the fetal, placental, and uterine wall tissues of rhesus macaques. Third, this study focused exclusively on evaluating the existence of bacterial communities in the fetal and placental tissues of rhesus macaques. The existence of eukaryotic microbial communities and viruses in these tissues was not considered.

### Conclusions

Using bacterial culture, 16S rRNA gene qPCR, and 16S rRNA gene sequencing, there was not consistent evidence of bacterial communities inhabiting the fetal and placental tissues of rhesus macaques. This study provides further evidence against the *in utero* colonization hypothesis and the existence of a placental microbiota. If there are intrauterine bacterial communities, they are limited to the uterine wall.

## Acknowledgements

We gratefully acknowledge Sarah Davis and Christy Johnson of the California National Primate Research Center at the University of California Davis for critical logistical support and for monitoring and reporting the results of bacterial cultures, respectively.

## MATERIALS AND METHODS

### Study subjects and sample collection

This was a cross-sectional study of four rhesus macaque dams undergoing cesarean delivery of a ∼130-day (129-132) gestational age fetus without labor. These dams were among the saline control subjects of a broader study at the California National Primate Research Center within the University of California Davis, with approved procedures and protocols through IACUC #20330 in 2018. Upon delivery of the fetus, a uterine wall biopsy and Dacron swabs (Medical Packaging Corp., Camarillo, CA) of the uterine wall decidua and the myometrium were collected (these uterine wall swabs were not collected from subject 3). Dams did not receive antibiotics, including intraoperative prophylaxis, prior to sampling.

The placenta and umbilical cord were placed in an autoclave-sterilized container and covered. Rhesus macaque fetuses were euthanized with pentobarbital (100 mg/kg) prior to necropsy. The fetal liver, heart, and brain were snap frozen in sterile 50 ml conical tubes. The fetal colon was also placed in a sterile 50 ml conical tube and it, along with the placenta and umbilical cord, were immediately transported to a biological safety cabinet in a nearby laboratory within the National Primate Research Center for further processing.

Study personnel donned sterile surgical gowns, masks, full hoods, and powder-free exam gloves during sample processing. Sterile disposable scissors and forceps were used throughout, and new scissors and forceps were used for each organ and each specific organ site that was sampled. Dacron swabs and ESwabs (BD Diagnostics, MD) were collected for molecular microbiology and bacterial culture, respectively.

For the placenta, samples were collected midway between the longest distance from the cord insertion point to the edge of the placental disc. Dacron swabs were collected from three sites of the chorionic plate (top of the amnion, amnion-chorion interface, and subchorion) and from the villous tree and basal plate. Eswabs were collected from two sites on the chorionic plate (amnion-chorion interface and subchorion) and from the villous tree. From a separate section of the placental disc, distant from the area where swabs were taken, a full-thickness (i.e. chorionic plate through to basal plate) portion (∼1 cm^2^) of the placenta was collected. A cross-section of the umbilical cord was also collected. The fetal colon was sectioned into proximal, central, and distal portions. The proximal and distal portions of the colon were sliced open lengthwise and the luminal contents and mucosal lining were swabbed with Dacron swabs and ESwabs. Dacron swabs and tissues were frozen at −80°C. ESwabs were processed for culture.

### Bacterial culture

Within three hours of fetal delivery, ESwab samples for bacterial culture were processed in a biological safety cabinet by study personnel wearing a sterile surgical gown, mask, full hood, and powder-free exam gloves. Specifically, ESwab buffer solutions were added to SP4 broth with urea (Hardy Diagnostics, Santa Maria, CA) and were plated on blood agar (trypticase soy agar with 5% sheep blood) and chocolate agar. Samples of the chorionic plate (amnion-chorion interface and the subchorion), villous tree, and fetal distal colon were inoculated on each culture medium. ESwab samples of the fetal proximal colon were inoculated on blood and chocolate agar, but not SP4 broth. Blood and chocolate agar plates were incubated under aerobic (5% CO_2_) and anaerobic (BD GasPak EZ anaerobic pouch; Franklin Lakes, NJ) atmospheres at 37°C for seven days. SP4 broth was only incubated under aerobic conditions. Negative and positive (blood and chocolate agar inoculated with a human buccal ESwab) culture media controls were incubated alongside the rhesus macaque samples for seven days.

### Taxonomic identification of individual bacterial isolates

Bacterial isolates (i.e. colonies) recovered from rhesus ESwab samples were taxonomically identified based upon their 16S rRNA gene sequence identity. The 16S rRNA gene of each bacterial isolate was amplified using the 27F/1492R primer set (5’-AGAGTTTGATCMTGGCTCAG-3’/5’-TACCTTGTTACGACTT-3’) and then bi-directionally Sanger sequenced by GENEWIZ (South Plainfield, NJ) using the 515F/806R primer set (5’-GTGYCAGCMGCCGCGGTAA-3’/5’-GGACTACNVGGGTWTCTAAT-3’), which targets the V4 hypervariable region of the 16S rRNA gene. Forward and reverse reads were trimmed using DNA Baser software (http://www.dnabaser.com/) with default settings, and assembled using the CAP (contig assembly program) of BioEdit software (v7.0.5.3; Carlsbad, CA), also with default settings. The taxonomic identities of individual bacterial isolates were determined using the Basic Local Alignment Search Tool (BLAST) (103) with a percent nucleotide identity cutoff of 100%.

### DNA extraction from swab and tissue samples

All Dacron swab and tissue samples were stored at −80° C until genomic DNA extractions were performed. These extractions were performed in a biological safety cabinet by study personnel wearing sterile surgical gowns, masks, full hoods, and powder-free exam gloves. DNA was extracted from swab and tissue samples separately, and the order of extractions was randomized within each sample type (i.e. swabs and tissues).

For DNA extraction from Dacron swabs, a DNeasy PowerLyzer PowerSoil kit (Qiagen, Germantwon, MD) was used with minor modifications to the manufacturer’s protocol. Specifically, after UV sterilizing all kit reagents (excluding the spin column), 500 μl of bead solution, 200 μl of phenol:chloroform:isoamyl alcohol (pH 7–8), and a swab were added to the supplied bead tube. The tube was inverted and, after a 10-minute incubation at room temperature, the tube was vortexed and centrifuged, and the swab was removed. Sixty μl of Solution C1 were added to the tube prior to bead beating two times at 30 seconds. The remainder of the DNA extraction process was as previously published (84).

For DNA extraction from tissues, a Qiagen PowerSoil DNA Isolation kit was used. Minor modifications from the manufacturer’s protocol were that all kit reagents (excluding the spin column) were UV-sterilized, cells within samples were lysed by mechanical disruption three times for 30 seconds using a bead beater, and DNA was eluted from the spin column using 60 μl of C6 solution. For these extractions, 0.140 – 0.200 grams of tissue were used. For fetal heart and liver samples, longitudinal sections were taken from the middle of specimens. For umbilical cord samples, transverse sections were taken. Purified DNA was stored at −20° C.

### 16S rRNA gene quantitative real-time PCR (qPCR)

#### Preliminary inhibition test

A preliminary test was performed to determine whether DNA amplification inhibition existed among the different sample types (tissues and swabs by body site). Purified DNA from each sample was first quantified using a Qubit 3.0 fluorometer with a Qubit dsDNA Assay kit (Life Technologies, Carlsbad, CA). For the inhibition test, 2.0 μl of purified *Escherichia coli* ATCC 25922 (GenBank accession: CP009072) genomic DNA containing seven 16S rDNA copies per genome was spiked into 4.0 μl of purified DNA from samples (normalized to 80 ng/μl genomic DNA when possible), which had been serially diluted with Qiagen Solution C6 by a factor of 1:3. Two μl of each spiked sample were then used as a template for qPCR. All reactions in each qPCR run were spiked with an equal amount of DNA (either 3.28 × 10^3^ or 5.92 × 10^3^ 16S rRNA gene copies).

Total bacterial DNA abundance within spiked samples was measured via amplification of the V1 - V2 region of the 16S rRNA gene according to the protocol of Dickson et al (104), with previously published minor modifications (84). Raw amplification data were normalized to the ROX passive reference dye and analyzed using the Thermo Fisher Cloud and Standard Curve (SR) 3.3.0-SR2-build15 with automatic threshold and baseline settings. Cycle of quantification (Cq) values were calculated for samples based on the mean number of cycles required for normalized fluorescence to exponentially increase.

The inhibition test indicated a low level of inhibition for most rhesus macaque tissue and swab DNA sample types. Therefore, all tissue, swab, blank technical control, and positive control DNA template samples were diluted with Qiagen Solution C6 by a factor of 1:4.5 prior to qPCR. The positive controls were six human urine samples: three urine samples (genomic DNA from 10 ml urine) were run alongside the rhesus tissue samples, and three different urine samples (genomic DNA from 1 ml urine) were run alongside the rhesus swab samples. The collection of urine samples and their use for research was approved by the Human Investigation Committee of Wayne State University and the Institutional Review Board of the Eunice Kennedy Shriver National Institute of Child Health and Human Development. All subjects provided written informed consent for participation.

#### qPCR data generation

Total bacterial DNA abundance within rhesus macaque samples was measured by qPCR as described above for the inhibition test, with each sample being tested individually across triplicate runs. To estimate qPCR efficiency based on the slope of a standard curve and to determine the concentration of 16S rRNA gene copies in samples, a standard curve containing seven 10-fold serial dilutions (three replicates each) ranging from either 9.52 × 10^6^ to 10.0 16S rRNA gene copies (tissue samples) or 9.97 × 10^6^ to 10.0 16S rRNA gene copies (swab samples) was included in each run. All individual qPCR reactions had an efficiency ≥ 92.04% (slope ≤ - 3.5287).

### 16S rRNA gene sequencing of swab and tissue sample extracts

Amplification and sequencing of the V4 region of the 16S rRNA gene was performed at the University of Michigan’s Center for Microbial Systems as previously described (30, 84), except that library builds were performed in duplicate using 40 cycles of PCR and pooled for each individual sample prior to the equimolar pooling of all sample libraries for multiplex sequencing. Three human urine (1 ml) samples were included as positive controls. Sample-specific MiSeq run files have been deposited on the NCBI Sequence Read Archive (BioProject ID PRJNA610218).

Raw sequence reads were processed using DADA2 (v 1.12) (105). An analysis of 16S rRNA gene amplicon sequence variants (ASVs), defined by 100% sequence similarity, was performed using DADA2 in R (v 3.5.1) (https://www.R-project.org), and the online MiSeq protocol (https://benjjneb.github.io/dada2/tutorial.html) with minor modifications. These modifications included allowing truncation lengths of 250 bp and 150 bp and a maximum number of expected errors of 2 bp and 7 bp for forward and reverse reads, respectively. To allow for increased power to detect rare variants, sample inference allowed for pooling of samples. Additionally, samples in the resulting sequence table were pooled prior to removal of chimeric sequences. Sequences were then classified using the “silva_nr_v132_train_set” database with a minimum bootstrap value of 80%, and sequences that were derived from Archaea, Chloroplast, or Eukaryota were removed.

### Statistical analysis

Sample bacterial loads were assessed through cycle of quantification values obtained from qPCR. Differences in bacterial loads between fetal, placental, and uterine wall samples and background technical controls (i.e. sterile Dacron swabs and blank DNA extraction kits) were evaluated using Mann-Whitney U tests with sequential Bonferroni corrections applied.

Analyses of the 16S rRNA gene profiles of samples were limited to those with a minimum of 500 quality-filtered 16S rRNA gene sequences and a Good’s coverage ≥ 95.0% after uniform subsampling of all samples to 500 sequences. The average Good’s coverage values of swab and tissue samples after subsampling were 98.9 ± 0.8 SD and 99.1 ± 0.5 SD, respectively. Heat maps of the 16S rRNA gene profiles of samples were generated using heatmap.2 in the gplots library for R (version 3.5.1). The R package decontam (106) was utilized to identify ASVs that were potential background DNA contaminants using the “IsNotContaminant” method with a prevalence threshold of P = 0.5. The decontam analyses were run separately for the swab and tissue samples.

The alpha diversity of sample ASV profiles was characterized using the Chao1 index to address profile richness and the Shannon and Simpson (1 – D) indices to address profile heterogeneity. Differences in alpha diversity between rhesus macaque and background technical control samples were evaluated using Mann-Whitney U and t-tests with sequential Bonferroni corrections applied.

The beta diversity of ASV profiles among fetal, placental and uterine wall samples and background technical controls was characterized using the Bray-Curtis similarity index. Bray-Curtis similarities in sample ASV profiles were visualized using Principal Coordinates Analysis (PCoA) plots and statistically evaluated using non-parametric multivariate ANOVA (NPMANOVA). PCoA plots were generated using the vegan package (version 2.5.5) in R. All statistical analyses were completed using PAST software (v 3.25) (107).

